# Predicting cyanobacterial oxic diazotrophy and prioritizing candidate protein families beyond known Nif proteins

**DOI:** 10.64898/2026.10.11.758377

**Authors:** James Young, Shiva Aryal, Dillon Nelson, Dheeraj Raya, Etienne Z. Gnimpieba, Ruanbao Zhou

## Abstract

**Background:** Diazotrophic cyanobacteria must reconcile oxygenic photosynthesis with the oxygen sensitivity of nitrogenase. Engineering studies have demonstrated nitrogenase activity in a non-diazotrophic cyanobacterium while highlighting the importance of accessory functions and oxygen management. Robust aerobic nitrogen fixation remains challenging, motivating systematic identification of candidate support functions beyond the canonical Nif machinery.

**Results:** We modeled *nifHDK*-defined diazotrophic potential from protein-family presence or absence in 426 cyanobacterial assemblies. Four classifiers evaluated with genus-blocked cross-validation achieved mean ROC AUCs of 0.853 to 0.893, with purity and genus-breadth filters fitted within training folds. Restricting the panel to 401 assemblies with current NCBI status gave AUCs of 0.806 to 0.881. Family-content logistic regression outperformed a genome-size-only baseline across ten genus partitions. Two inventories describe candidates across the full panel: 981 high-purity families and 473 families with positive residual associations (accompanied by three ubiquitous controls). Model importance orders the residual-association inventory. A separate heuristic score combines inventory membership with proteomics, condensate-driver ranks and genus-based morphotype proxies to prioritize 167 families for experiments. Proteomics annotations tracked detection opportunity; the condensate contrast between inventories attenuated after prevalence adjustment. These annotations help choose experiments but do not independently validate accessory functions.

**Conclusions:** Protein-family content predicts *nifHDK*-defined diazotrophic potential, with performance sensitive to assembly inclusion, taxonomy, genome size and taxonomic weighting. The two candidate inventories distinguish high carrier purity from residual association, providing complementary resources for prioritizing previously uncharacterized functions associated with cyanobacterial oxic diazotrophy.

## Background

Industrial ammonia synthesis has reshaped global agriculture and the nitrogen cycle [1]. Biological nitrogen fixation converts atmospheric dinitrogen into bioavailable ammonia through nitrogenase, an oxygen-sensitive metalloenzyme [2]. This oxygen sensitivity creates a central physiological contradiction for oxygenic phototrophs. The cells best positioned to sustainably power nitrogen fixation through photosynthesis also produce the oxygen that damages the enzyme. Cyanobacteria resolve this conflict through multiple documented strategies, including heterocyst differentiation in filamentous taxa [3], respiratory protection [4], electron-transfer remodeling, hydrogen recycling [5], and specialized regulation of nitrogen metabolism. Prior work from unicellular systems shows that oxic, light-driven nitrogenase activity can occur under continuous illumination even with net-production of oxygen [6], reinforcing that diazotrophy in cyanobacteria may have yet-to-be elucidated capabilities in balancing or even integrating photosynthesis and nitrogen fixation.

Engineering nitrogen fixation into oxygenic phototrophs remains attractive for sustainable agriculture, biomanufacturing, and broader bio-solar nitrogen-economy concepts [7]. Engineering studies and reviews identify challenges to obtaining robust activity from transferred *nif* structural genes [8–11]. Accessory functions are required for cofactor assembly, electron supply, oxygen management, metal trafficking, redox balance, stress protection, and regulatory timing [12]. Many of these functions are dispersed outside canonical *nif* clusters, condition dependent, poorly annotated, or lineage specific. As a result, the field needs ranked candidate inventories beyond descriptive lists of genes found near *nifHDK*.

Comparative bioinformatics can identify gene or protein families enriched in diazotrophs. Microbial genotype-phenotype associations remain vulnerable to phylogenetic structure, horizontal transfer, and correlated accessory genomes [13–15]. A family may appear associated with diazotrophy because it is carried by one successful diazotrophic lineage, because it is physically linked to a transferred island, or because it contributes to a function that supports nitrogen fixation across lineages. We framed the problem as protein-family modeling. We tested whether protein-family content predicts *nifHDK* status across held-out genus labels and used full-panel family associations and model ranks to nominate hypotheses beyond a high-purity inventory.

We also compared protein-family rankings with a separate gene-level feature list. Overlap and category composition were treated as descriptive because the original gene-feature construction could not be independently reconstructed. Protein families connect genome content to proteomics and functional annotation, although clustering thresholds and mapping choices limit comparisons.

Here, we analyze a cyanobacterial protein-family atlas for *nifHDK*-defined diazotrophic potential, following the broader comparative-bioinformatics logic that accessory-family content can expose trait-associated patterns [16]. We evaluate prediction with genus-blocked classifiers and treat feature importance as descriptive model behavior. We separate high-carrier-purity families from a residual association inventory, compare protein-family and gene-level rankings, and annotate candidates to guide experiments.

## Results

### Cyanobacterial protein-family matrix

The primary panel contains 426 cyanobacterial assemblies, including 112 *nifHDK*-positive and 314 comparison genomes across 75 stored genus labels (Fig. 1, Table 1). Its presence/absence matrix supplies the candidate inventories and downstream annotations.

**Figure 1.**
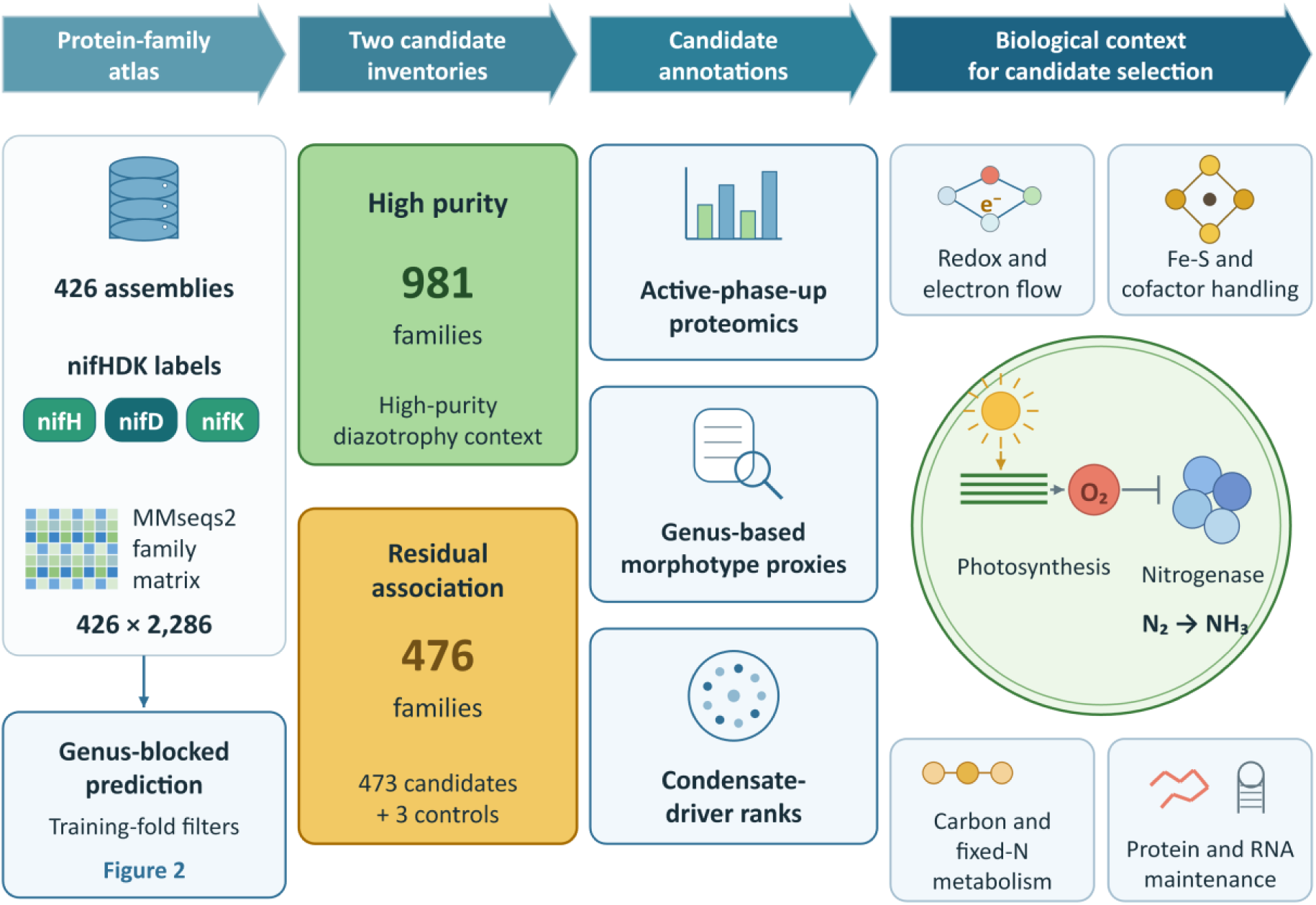
Study design and candidate annotation. The 426-assembly panel contains 112 *nifHDK*-positive genomes and 75 stored genus labels. MMseqs2 clustering at 80% identity and coverage, followed by a 40-genome prevalence filter, produced a 426 × 2,286 presence/absence matrix. Full-panel analysis identified 981 high-purity families and a 476-family residual-association inventory comprising 473 positive carrier/non-carrier contrasts and three ubiquitous controls. Model importance orders the residual inventory without imposing an inclusion threshold. A separate genus-blocked prediction analysis fits purity and genus-breadth filters within each training fold. Proteomics, morphotype proxies and condensate ranks annotate candidates for experimental selection. The prioritization score yields a 167-family shortlist after exclusions. The three ubiquitous controls and terms based on incompatible cross-atlas identifiers are excluded. Product-label comparison with the related atlas is descriptive and contributes no score. The biological contexts shown motivate experiments; they do not establish functional modules.

**Table 1.** Primary protein-family atlas and prediction analysis.

| Quantity | Primary atlas |
| --- | --- |
| Genome set | 426 assemblies from 489 screened records; RefSeq accession and completeness criteria |
| Feature representation | Protein-family presence/absence matrix |
| Protein-family matrix | 426 genomes × 2,286 families |
| Candidate inventories | 476 residual-association entries, including 3 ubiquitous controls; 981 high-purity families |
| Cross-validation design | Stored genus labels held within folds; purity and genus-breadth filters fitted within training folds |
| XGBoost mean ROC AUC ± fold SD | $0.893 \pm 0.078$ |

Predicted proteins were clustered into protein families with MMseqs2 [17] using 80% identity and 80% coverage settings. For supervised learning, families present in at least 40 genomes were retained, producing a 426-genome by 2,286-family binary matrix. Presence required at least one protein-family member in a genome. The models used family presence or absence.

ROC AUC is the unweighted mean ± sample standard deviation across five folds blocked by stored genus labels. Feature eligibility was determined within each training fold using a fixed family matrix. Candidate inventories and their importance ranking describe the full panel; cross-validation evaluates the prediction procedure.

### Genus-blocked classifiers recover diazotrophy signal from protein-family content

We trained four classifiers on the primary protein-family presence/absence matrix, including random forest [18], gradient boosting, logistic regression, and XGBoost [19]. Primary cross-validation kept each stored genus-name token within one fold. Purity and genus-breadth filters were fitted only to each training fold, using the fixed family matrix and stored genus assignments. The repository methods note documents the earlier use of a global purity filter and its replacement in prediction evaluation. This prevents held-out labels from selecting model features but does not remove all phylogenetic dependence.

With training-fold purity and genus-breadth filtering, mean ROC AUC was 0.853 ± 0.181 for random forest, 0.868 ± 0.124 for gradient boosting, 0.878 ± 0.095 for logistic regression, and 0.893 ± 0.078 for XGBoost (Fig. 2, Table 1). Mean average precision was 0.805, 0.790, 0.797 and 0.847, respectively. These estimates assess prediction from the residual feature space. They do not validate individual families in the full-panel ranking.

**Figure 2.**
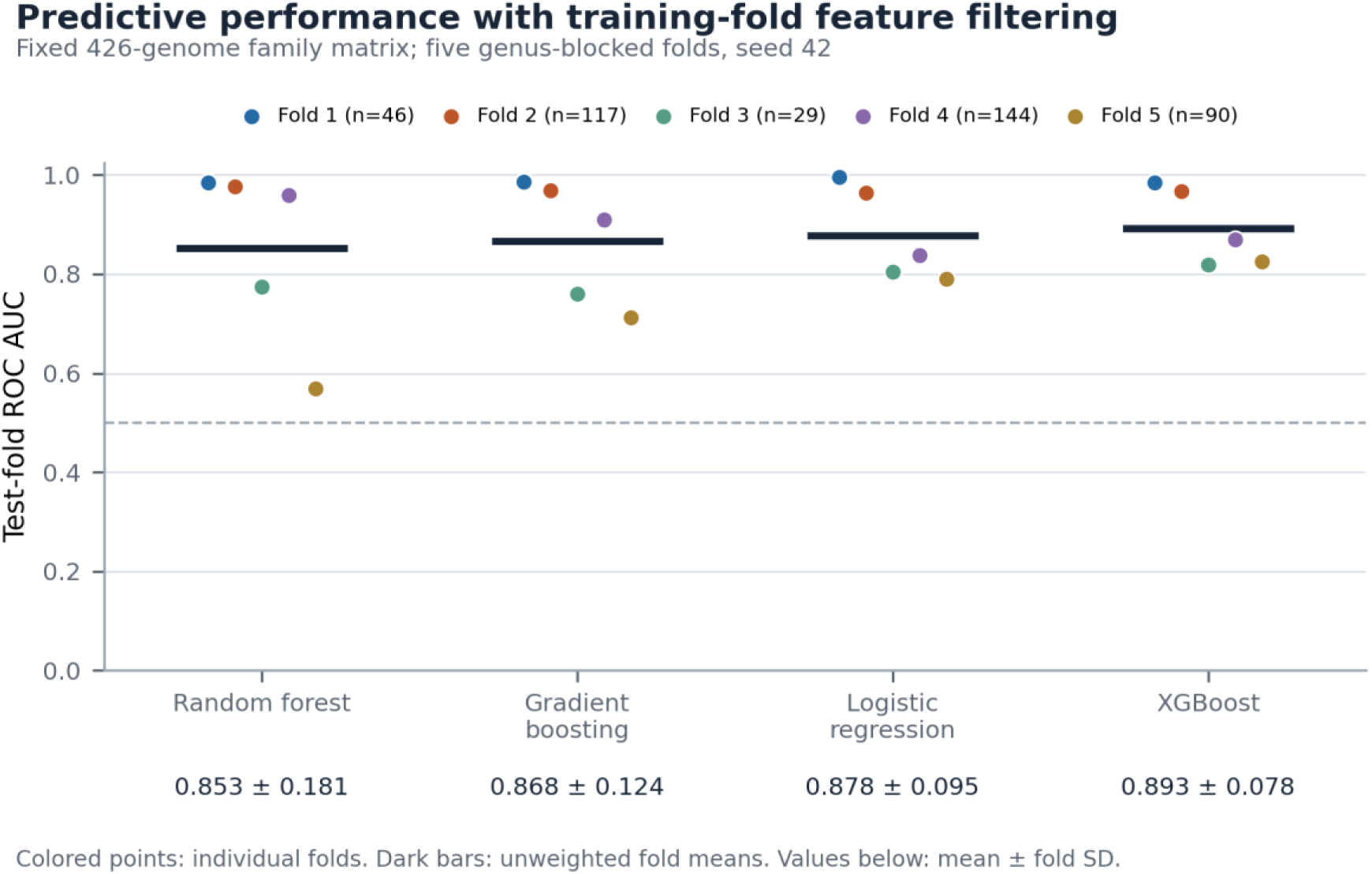
Predictive performance with training-fold feature filtering. Each point is test-fold ROC AUC after excluding families with training-carrier diazotroph fractions ≤0.10 or ≥0.90 and families represented in fewer than three training genera. Colored points identify the same five folds blocked by stored genus labels for all models; bars and printed values show unweighted fold means and sample standard deviations. The starting 2,286-family matrix was constructed across the full genome panel. These results evaluate the residual-feature procedure on that fixed matrix; they do not independently validate the full-panel candidate ranking. Fold sizes range from 29 to 144 genomes. Sample standard deviation describes variation across folds and is not a confidence interval. Size-only and genus-weighted sensitivity summaries accompany the prediction outputs.

Performance depended on the unit of aggregation. Across the four models, genus-weighted fold-mean AUC was 0.842–0.871, compared with 0.853–0.893 using equal genome weights within each fold. Pooling all out-of-fold predictions gave AUC 0.856–0.921 with equal genome weights and 0.782–0.815 when each genus received equal total weight. Pooled values also compare scores from different fitted models and are reported as sensitivity summaries, not interchangeable estimates. A logistic regression using assembly length alone reached mean fold AUC 0.816 ± 0.131; one using the number of retained families reached 0.749 ± 0.216. Broad genome properties therefore contribute substantial predictive information, and family-content discrimination alone does not isolate accessory mechanisms.

Across ten genus partitions (seeds 42–51), family-content logistic regression outperformed assembly-length-only logistic regression in every partition. Its fold-mean AUC ranged from 0.800 to 0.916, versus 0.726–0.875 for assembly length; the paired AUC difference ranged from 0.018 to 0.147. Adding assembly length to the family model changed mean AUC by less than 0.002 in either direction across these partitions. Thus, family content contains predictive information beyond these simple size summaries, while the result remains sensitive to the partition and does not identify causal family effects. These overlapping partitions are a sensitivity analysis, not ten independent experiments.

An NCBI status check on 30 September 2026 classified the 426 archived assembly versions as 401 current, 22 suppressed and three previous; six suppressed assemblies were *nifHDK*-positive. Restricting analysis to current versions and refitting models within the original folds yielded mean ROC AUCs of 0.806–0.881, compared with 0.853–0.893 for the full panel. Random forest showed the largest decline, from 0.853 to 0.806. These analyses retained the original family vocabulary. Separately, MIT S9506 and SS51 had identical retained profiles across folds, without establishing whole-genome identity. A broader conservative grouping analysis yielded AUCs of 0.820–0.852; its effects cannot be attributed solely to that pair. Complete cohort, taxonomy and grouping results are documented in the accompanying Cohort and taxonomy sensitivity analysis note.

The four classifiers assess how prediction depends on the modeling approach. Tree-based models can represent nonlinear combinations, while logistic regression provides an additive comparison. Their performance does not establish a biological interaction mechanism or show that individual ranked families improve prediction beyond correlated alternatives. The residual-association inventory is defined by a univariate association rule; model importance orders its members.

### Two non-overlapping inventories separate high carrier purity from residual association

The strongest single-family diazotrophy markers include core Nif machinery and other near-diagnostic families. If these families enter the model unconstrained, they can dominate classification and obscure the accessory context most useful for engineering. We therefore separated high-purity families from the residual feature space.

The high-purity inventory contains 981 families for which at least 90% of carrier genomes are labeled diazotrophs. These families were excluded from the model feature ranking. Known nitrogenase and Nif machinery families provide expected biological reference points. Other product annotations include membrane stress, transport, regulation, respiration, redox balance, and metallocluster or Fe-S biology. Carrier purity is the fraction of a family’s carriers labeled diazotrophs. It does not measure the fraction of diazotroph genomes carrying that family, so high purity can coexist with limited diazotroph coverage.

The residual-association inventory contains 473 families with a positive difference in diazotroph frequency between carriers and non-carriers. Three ubiquitous families accompany these candidates as controls, giving 476 inventory entries. Because those controls have no non-carriers, their association cannot be estimated; the source implementation included them by substituting zero for the empty comparison group. All 476 families appear in the four model-importance tables. Their mean importance percentiles determine their order, without a nonzero-importance or family-level significance threshold. The controls have zero importance in all four models and are excluded from the experimental shortlist. Inventory membership therefore identifies candidates for follow-up rather than individually validated predictors.

The candidate inventories and their importance ranking describe the full panel. Cross-validation determines feature eligibility separately within each training fold, retaining 568–653 families, and does not redefine the inventories. The repository methods note documents the original filtering implementation and its boundary checks.

### Residual associations identify named accessory-function candidates

Product-name matching classified 74 of 476 residual-association inventory entries (16%) as generic or hypothetical. These labels describe annotation specificity, not biological novelty. The repository annotation note defines the keyword categories and interpretive labels.

The residual-association inventory includes mechanistically suggestive accessory functions as well as signals that may partly reflect lineage structure. Candidate annotations include nitrogen assimilation through GOGAT; Fe-S and metal homeostasis through SufB, Fe-S accessory proteins, and CobW; tetrapyrrole and oxygen physiology through Mg-protoporphyrin cyclase and the DPOR pair; respiration and electron flow through NDH-1; carbon/reductant metabolism through transketolase and oxidative pentose-phosphate or TCA-linked enzymes; nitrogen regulation through NtcA and P-II; and proteostasis through DnaK and GroEL. These examples describe the inventory and motivate experiments; functional labels do not establish coordinated genomic modules or mechanistic requirements.

Interpretive labels refer to FOX (fixation in the presence of oxygen), the phenotype considered in the prior multi-omic model [20]. These labels provide biological context and contribute no score.

For the inventory ranking, we converted absolute importances in each model’s summary table to percentiles and averaged the percentiles. These tables summarize fold fits after feature filtering across the full panel. The prioritization score instead assigns a fixed weight to inventory membership and combines it with proteomics, condensate and morphotype annotations. Figure 3 displays all 22 Tier A families. Recomputing importance with training-fold feature filtering changed the four-model consensus ordering: Spearman ρ was 0.831 across the 476 inventory entries, and 33 of the full-panel top 50 remained in the top 50. Both rankings are provided. Neither rank agreement nor the annotation score independently validates a candidate.

**Figure 3.**
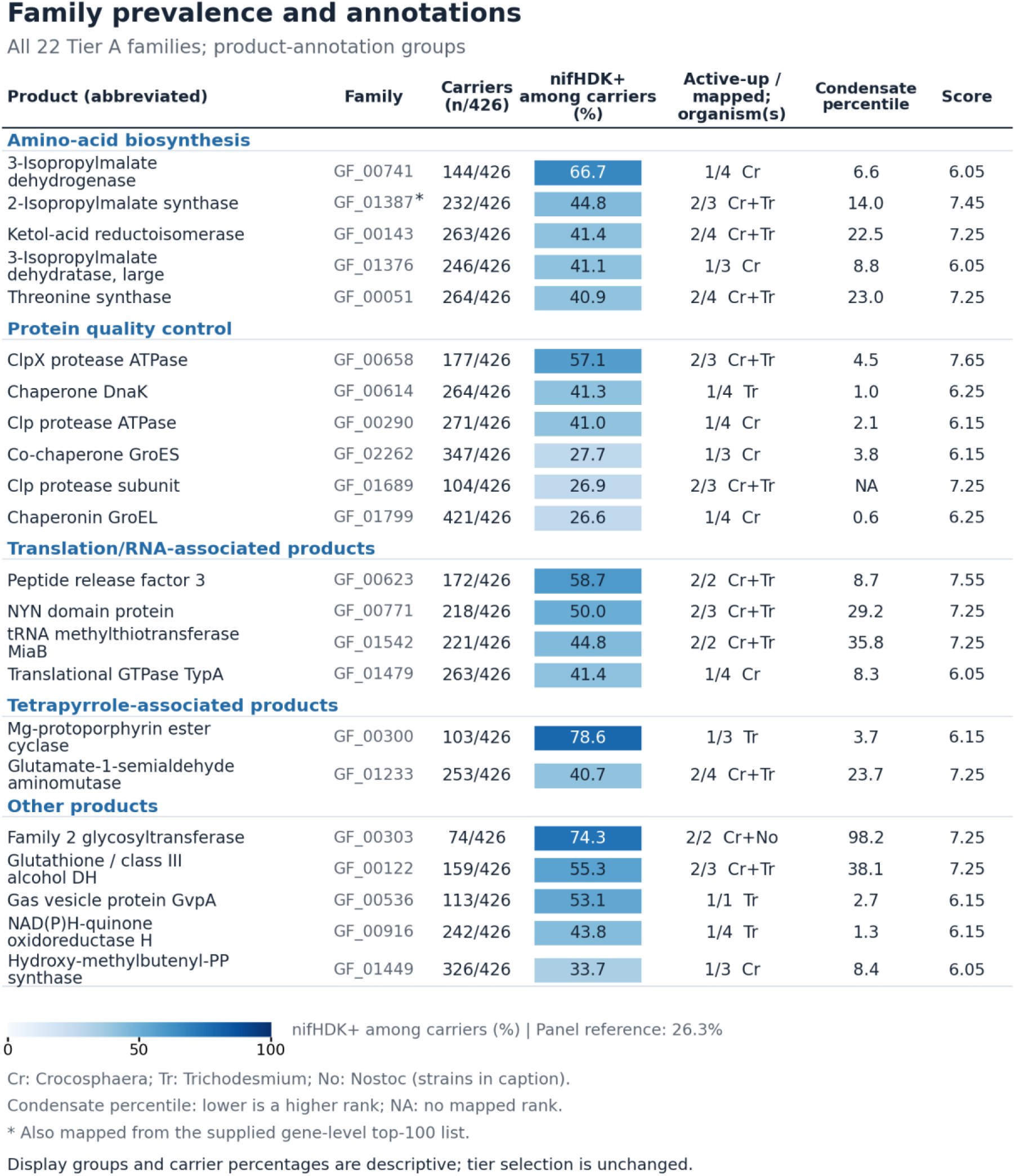
Family prevalence and annotations for all 22 Tier A residual-association families. Rows use heuristic product-annotation groups and, within each group, decreasing carrier purity followed by family ID. These display groups do not change scores or tier membership. Carriers gives the number of panel genomes containing the family. Shading shows the percentage of those carriers labeled *nifHDK*-positive, not coverage of positive genomes or evidence strength; 112 of 426 panel genomes are positive (26.3%). Active-up gives studies with an active-phase-up annotation over studies with a mapped protein, followed by the contributing organisms: Cr, *Crocosphaera subtropica* ATCC 51142; Tr, *Trichodesmium erythraeum* IMS101; No, *Nostoc punctiforme* PCC 73102. Study coverage and response rules differ, so these are descriptive counts. Condensate percentile summarizes the best mapped protein per family; lower values indicate higher ranks. NA means no mapped rank. The prioritization score selected Tier A at ≥6.0 after the stated exclusions and is shown for reference. An asterisk marks a match in the supplied gene-level top-100 mapping. Product labels are abbreviated (DH, dehydrogenase; PP, diphosphate); full labels accompany the source data. These annotations guide experiments without independently validating candidates or establishing genetic requirement.

### High-purity families include core diazotrophy machinery and accessory candidates

The high-purity inventory includes annotated nitrogenase structural and cofactor/assembly families, alongside non-Nif candidates and less specific product labels.

The Nif machinery families provide expected reference components. High-purity non-Nif families form a context inventory: their carriers are predominantly diazotroph-labeled, but this does not establish that the families are required by diazotrophs or by an engineered recipient. Some may be lineage-linked passengers or markers of transferred islands; others may encode useful support functions. Functional annotation, carrier distribution and physiological measurements can guide candidate selection, while perturbation and recipient-specific tests are needed to establish necessity or sufficiency.

### Morphotype proxies describe carrier representation

Because cyanobacterial diazotrophy is distributed across unicellular and filamentous strategies, we asked which candidate families are represented in both morphotype contexts. The current metadata lack a curated morphology field, so this analysis used genus-based proxies. A strict unicellular proxy contained 15 diazotroph genomes across 9 stored genus labels, while a broader proxy added *Chroococcidiopsis*, *Cyanobacterium*, and *Pleurocapsa*, yielding 20 diazotroph genomes across 12 stored labels. All other diazotroph-labeled genomes were treated as a filamentous or non-unicellular proxy for this sensitivity analysis.

The primary morphotype-bridge flag requires at least one broad-proxy unicellular diazotroph and at least one filamentous-proxy diazotroph. This retained 420 of 476 residual-association and 188 of 981 high-purity families (Table 2). A stricter flag requiring at least three strict-proxy unicellular carriers and at least one filamentous-proxy carrier retained 210 residual-association and 6 high-purity families. These counts describe carrier representation under genus-based proxies. The inventories differ in prevalence, so this contrast does not by itself establish greater physiological breadth or portability of residual-association functions.

**Table 2.** Genus-based morphotype representation.

| Representation criterion | Definition | Residual-association inventory | High-purity inventory | Gene-level top-100 overlap |
| --- | --- | --- | --- | --- |
| Primary cross-morphotype bridge | $\geq 1$ broad-proxy unicellular diazotroph and $\geq 1$ filamentous-proxy diazotroph | 420/476 (88.2%) | 188/981 (19.2%) | 6 residual-association, 1 high-purity |
| Strict cross-morphotype bridge | $\geq 1$ strict-proxy unicellular diazotroph and $\geq 1$ filamentous-proxy diazotroph | 299/476 (62.8%) | 18/981 (1.8%) | 4 residual-association, 0 high-purity |
| Broad unicellular-breadth flag | $\geq 3$ broad-proxy unicellular diazotroph genomes and $\geq 1$ filamentous-proxy diazotroph | 289/476 (60.7%) | 12/981 (1.2%) | 4 residual-association, 1 high-purity |
| Strict unicellular-breadth flag | $\geq 3$ strict-proxy unicellular diazotroph genomes and $\geq 1$ filamentous-proxy diazotroph | 210/476 (44.1%) | 6/981 (0.6%) | 4 residual-association, 0 high-purity |
| *nif*/nitrogenase keyword set | Fixed 12-family keyword set, separate from morphotype criteria | 1 family | 11 families | Separate from the top-100 filter |

A separate 12-family *nif*/nitrogenase keyword set defines reference annotations and scoring exclusions. The NifD-like GF_00526 and NifH-like GF_00168 families pass the broad morphotype bridge, whereas the NifK-like GF_00158 family lacks broad-proxy unicellular carriers in this matrix. These differences warrant inspection of cluster membership and sequence coverage before inferring absence of the corresponding function. The set also contains annotated cofactor and support proteins and Nif3-like GF_00097. A keyword match alone does not establish a canonical nitrogenase role. Membership and scoring exclusions are documented in the repository.

The gene-driven top-100 comparison provided a representation-level stress test for the morphotype grading. Under the primary bridge flag, 6 residual-association families and 1 high-purity family overlap the gene-driven top-100 mapping. Under the stricter >=3 strict-unicellular flag, overlap drops to 4 residual-association families and 0 high-purity families. The overlapping strict-breadth families are GF_00488, GF_01387, GF_01690, and GF_02172, annotated as dihydroxy-acid dehydratase, 2-isopropylmalate synthase, CTP synthase, and GTP cyclohydrolase I FolE. These hits point to metabolism and cofactor-related functions shared across approaches. The low overall overlap reinforces that gene-level and protein-family approaches recover different biological views.

The primary morphotype bridge uses the broader unicellular proxy because the metadata contain no curated morphology column. Stricter criteria describe additional levels of carrier representation.

### Protein-family and gene-level representations produce different candidate rankings

A separate gene-level model used genus-blocked random forests on gene-level features and reported a top-100 list. To compare the two representations, we mapped the gene-level features into our protein-family space using all genes available for each gene-driven feature family. The mapping identified 23 protein families in the candidate inventories (7 residual-association and 16 high-purity) linked to the gene-driven features (Figure 4). This establishes partial concordance. Some gene-level signal recovers diazotrophy-associated protein-family context, especially the high-purity context set. The two approaches produce different top-ranked candidate sets; the degree of overlap also depends on successful cross-representation mapping.

**Figure 4.**
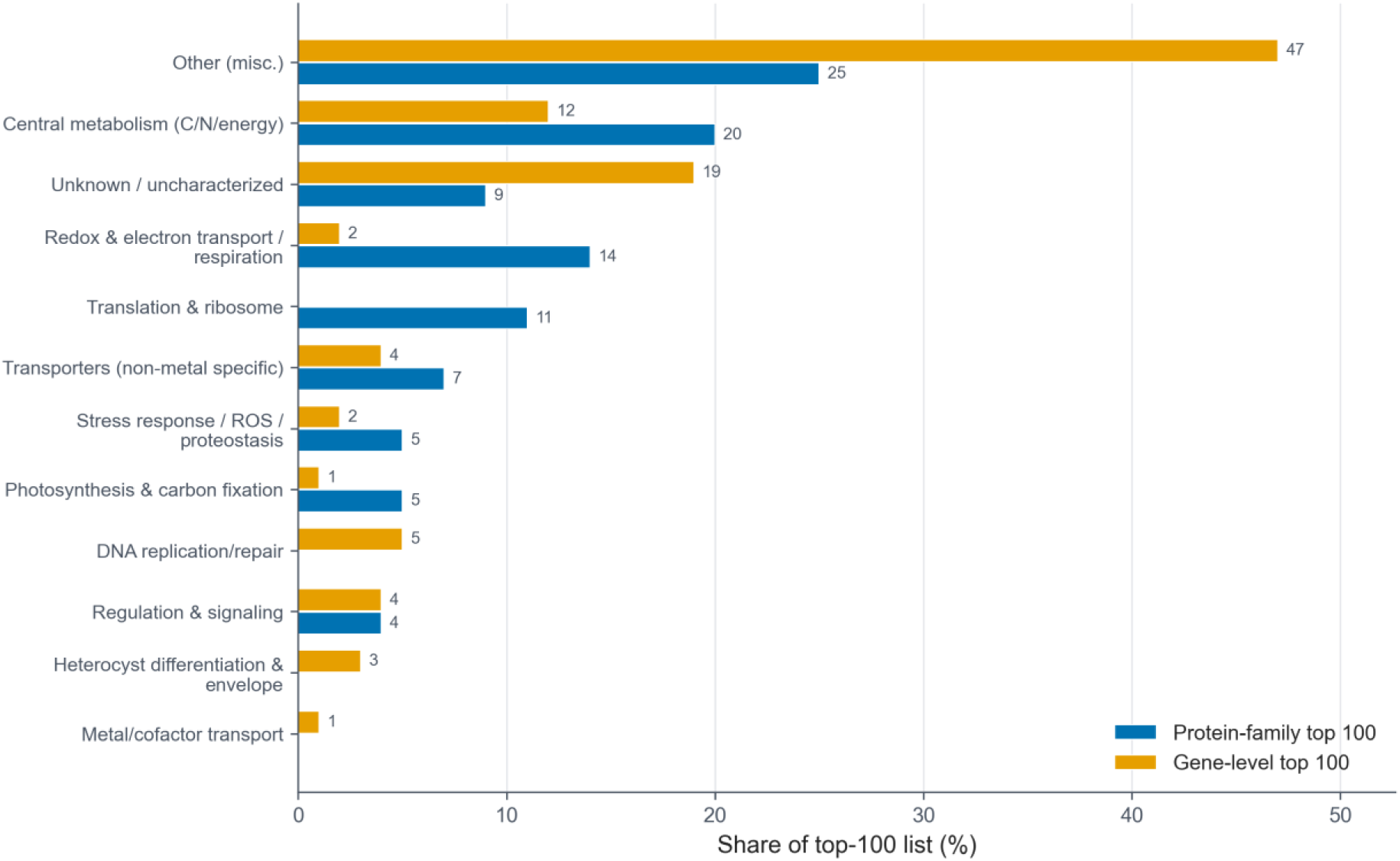
Descriptive comparison of protein-family and gene-level feature lists. The original keyword-based classification summarizes functional composition within two top-100 lists. The item-level table and plotting source reproduce these counts. Different representations produce different category profiles, but annotation quality and keyword rules affect this contrast; it does not establish greater biological relevance of either representation. The separate mapping comparison identifies 23 protein families in the candidate inventories linked to gene-driven features (7 residual-association and 16 high-purity), not necessarily 23 one-to-one feature matches. Stronger interpretability comparisons require consistent functional re-annotation of both lists.

The top-100 lists also differed under the current keyword annotations: 28 protein-driven features and 13 gene-driven features were assigned to diazotrophy-physiology categories. The gene-driven list contained 47% Other and 19% Unknown/uncharacterized, compared with 25% and 9% in the protein-driven list. Because these assignments depend on product-string quality and keyword coverage, they do not establish that one representation contains intrinsically more interpretable biology. The composition comparison remains descriptive pending consistent functional re-annotation of both lists.

Gene-level features may retain fine-grained locus and lineage variation, while protein families provide a convenient unit for comparison with proteomics and protein-function annotation. The present comparison supports treating these as complementary representations; it does not establish conserved functional islands or a causal advantage of either representation.

### A related protein-family atlas provides a product-level consistency check

We compared product annotations with the related analysis in the ERISE-BNERC repository Predictive Comparative Genomics for Diazotrophy (revision 85277c79). That analysis used 449 genomes, 2,551 protein-family features and a 503-family candidate set. It shared 416 assemblies with the primary panel, so the comparison tests consistency on largely shared data. Family identifiers differ between the clustering runs and cannot be treated as one-to-one biological units. The repository note Related-atlas comparison provides the versioned source links, reported prediction metrics and verification limits.

Normalized product/function labels provided an annotation-level comparison, without establishing one-to-one orthology. Of 503 related-atlas Tier 1 families, 442 had a product-label match in the current 1,457-family inventory, including 363 with informative labels after excluding generic hypothetical, uncharacterized, DUF-containing, domain-containing-protein and broad family-protein names. Of the 442 matched related-atlas families, 402 touched at least one current residual-association family and 153 touched at least one high-purity family; 113 touched both sets. These counts use related-atlas families as their unit and are not directly comparable with the gene-driven overlap counted in current protein families. Because 416 genomes are shared and matching uses product labels, this is a consistency check on largely shared data. It does not independently validate candidate functions and contributes no points to the composite score.

The prioritization score excludes terms transferred through raw family identifiers from the related atlas because those identifiers are incompatible. It also excludes the three ubiquitous controls from shortlist eligibility. With the stated weights and thresholds, the shortlist contains 22 Tier A, 140 Tier B and 5 high-purity-tier families (167 total); 65 have an active-phase-up annotation. The repository note documents the identifier mismatch and the resulting score and tier changes. Product-label agreement alone cannot substitute for a protein-membership or sequence-based family crosswalk.

### Proteomics timing and condensate-driver rankings annotate individual candidates

We annotated candidate families with study-specific proteomics responses relevant to nitrogen-fixing physiology: the dark nitrate-free contrast in *Crocosphaera subtropica* ATCC 51142, daytime abundance in *Trichodesmium erythraeum* IMS101, the late-light/early-dark D0/D3 window in *Cyanothece* sp. PCC 7822, and early heterocyst enrichment relative to filaments in *Nostoc punctiforme*. The operational active-up rules differ by source and do not all require a per-protein significance test. Active-phase-down, transition-window and non-target-phase responses remain annotations without adding score [21–24].

Across the two inventories, 139 of 1,457 families had at least one external active-phase-up proteomics annotation. Examples span protein maintenance (ClpX GF_00658, DnaK GF_00614 and GroEL GF_01799), RNA-associated functions (NYN-domain GF_00771 and MiaB GF_01542), nitrogen regulation (NtcA GF_00093 and P-II GF_01395), and carbon/redox metabolism (glucose-6-phosphate dehydrogenase GF_01818 and isocitrate dehydrogenase GF_00624). These annotations identify contexts for experiments without establishing enrichment beyond detection opportunity or a requirement for diazotrophy. The count includes the ubiquitous cytochrome b6 control GF_01297, which is excluded from the experimental shortlist.

The condensate-driver ranking supplied an exploratory sequence-based annotation from the resource described in the Nitrosomes preprint [25]. Of 31,021 ranked proteins, 2,125 mapped to the current pangenome crosswalk, representing 1,311 families: 369 residual-association, 934 high-purity, and 8 outside both inventories. The descriptive top-10% contrast was 64 of 369 residual-association families versus 67 of the 942 other mapped families (odds ratio 2.74). This comparison was not prevalence matched. Among mapped candidate families, the two inventories had median carrier counts of 117 and 55 genomes, respectively, and the inventory contrast attenuated after prevalence adjustment *as* reported below. We therefore retain condensate-driver rank as a candidate annotation rather than evidence of selective accessory biology.

High-ranked condensate candidates included GroEL GF_01799, DnaK GF_00614, NAD(P)H-quinone oxidoreductase subunit H GF_00916, magnesium-protoporphyrin IX monomethyl ester cyclase GF_00300, and 2-isopropylmalate synthase GF_01387. These rankings nominate sequence and localization questions for follow-up; they do not demonstrate condensate formation, recruitment, or a diazotrophy-specific function.

### Detection opportunity and prevalence constrain interpretation of the annotation layers

Among detected residual-association families (Fig. 5A), the odds of at least one active-phase-up annotation increased with the number of mapped proteomics studies (odds ratio 2.22 per additional study, 95% confidence interval 1.62–3.05; p = 8 × 10⁻⁷). After adjustment for that count and model rank, the strict unicellular-breadth flag had no detectable association with active-phase-up status in this model (odds ratio 0.89, 95% confidence interval 0.46–1.73; p = 0.73). The mean per-family fraction of mapped studies scored active-phase-up was 22.3% both with and without the strict breadth flag. Thus, repeated detection creates additional opportunities to accrue proteomics score, and the fraction of prioritized families carrying an active-phase-up annotation should not be read as independent convergence. Active-phase-up counts are also used to select the priority tiers themselves; the breadth regression does not directly test a detection-matched null for a shortlist selected partly on active-phase-up annotations.

**Figure 5.**
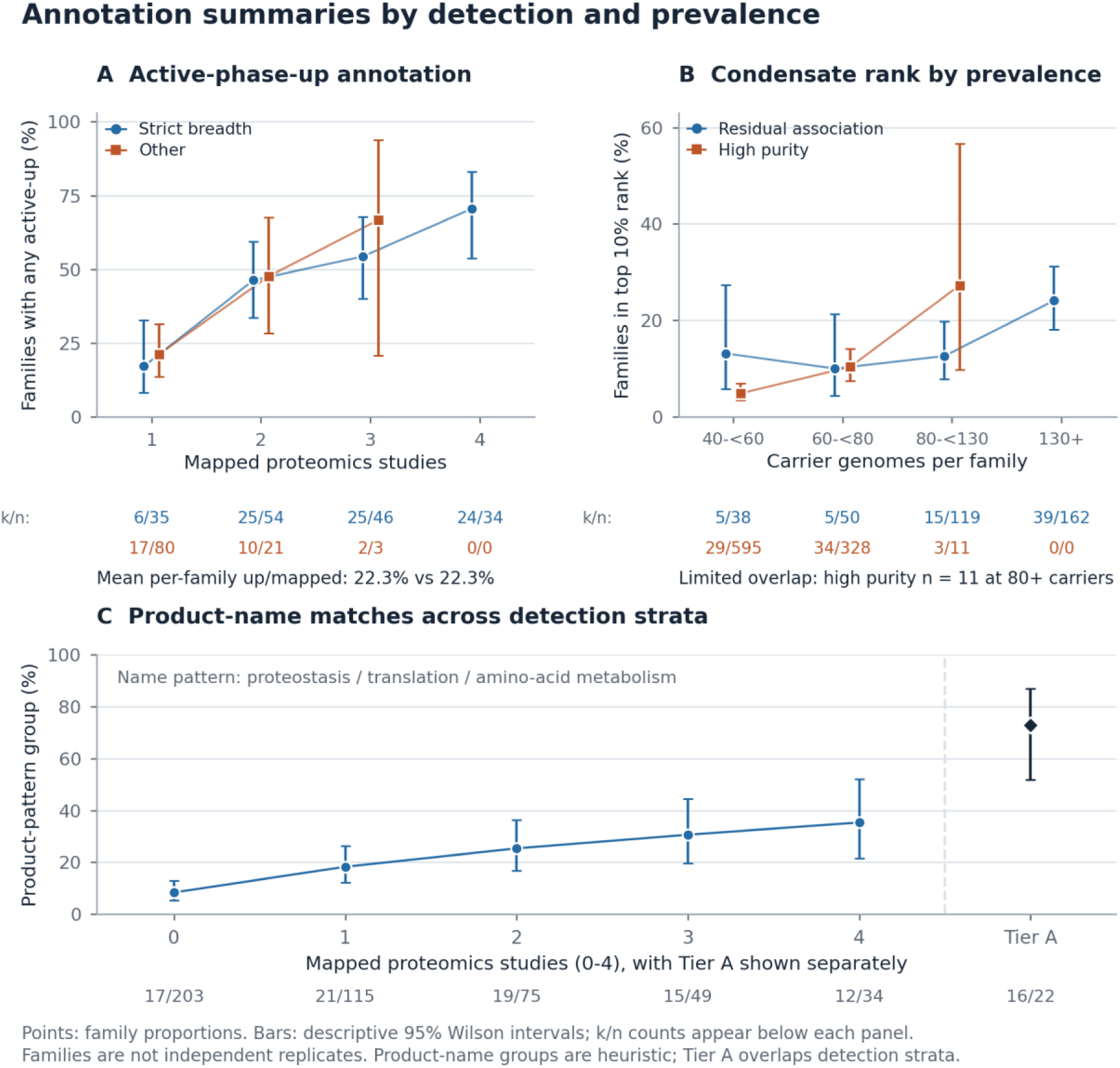
Detection opportunity and family prevalence constrain annotation summaries. (A) Among residual-association families mapped in at least one external proteomics study, the proportion with any active-phase-up annotation is shown by mapped-study count and the strict unicellular-breadth flag (>=3 strict-proxy unicellular and >=1 filamentous-proxy diazotroph carrier). The reported 22.3% values are means of each family’s active-up-study count divided by its mapped-study count, not pooled per-study probabilities. (B) The proportion of condensate-mapped candidate families in the top 10% of mapped family ranks is shown by inventory and carrier-genome count. Bins are half-open [40,60), [60,80), [80,130), and [130,infinity), matching the diagnostic stratification through 130 genomes. All nonempty strata are shown, including the 11 high-purity families at 80-129 carriers; no high-purity family has 130 or more carriers. (C) Matches to a product-name pattern covering proteostasis, translation, and amino-acid metabolism are shown across mapped-study counts in all 476 residual-association families. Diagnostic strata retain all 476 residual-association families, including three ubiquitous controls excluded from shortlist eligibility. Under the prioritization score, the pattern matches 16 of 22 Tier A families (72.7%). Tier A is a selected subset overlapping those strata, not an independent comparison group. Under each panel, k/n gives the numerator and denominator in matching series order and color; 0/0 denotes an empty stratum with no plotted estimate. Error bars are descriptive 95% Wilson intervals for family proportions and do not account for phylogenetic or other dependencies among families. Product-name matching is a heuristic diagnostic, not a validated functional classification. These patterns motivate detection- and prevalence-aware interpretation; they do not establish absence of biological roles for individual candidates.

For mapped candidate families, the residual-association versus high-purity odds ratio for top-10% condensate rank was 2.81 after adjustment for the number of mapped proteins per family (Fig. 5B). Adding carrier-genome count reduced that estimate to 0.97 (95% confidence interval 0.53–1.79; p = 0.93). Prevalence-stratified comparisons were inconsistent and included small strata. These diagnostics are consistent with prevalence dependence of the inventory contrast; they neither establish absence of condensate biology nor validate a prevalence-adjusted biological mechanism.

The prioritization score favors broadly distributed families by construction. Morphotype proxies supply 51.3% of the mean score for the 162 residual-association families in Tiers A and B, and all 22 Tier A families pass both scored breadth criteria. Of 140 Tier B families, 78 have neither scored proteomics nor condensate annotation. The score correlates weakly with full-panel model importance (Spearman ρ = 0.116 among the 473 estimable families; 0.104 across all 476 inventory entries). This is consistent with assigning a fixed weight to inventory membership. Tier labels describe heuristic score bands rather than independent agreement among evidence sources.

Product-name matches to the proteostasis, translation and amino-acid-metabolism pattern increased from 8% among families mapped in no proteomics study to 35% among those mapped in all four (Fig. 5C). This describes how product-name matches vary with study coverage. It does not establish a curated functional composition or explain the Tier A share by itself, because tier assignment also uses active-phase-up annotations. The repository annotation note reports the separate category co-occurrence analysis.

## Discussion

### Predictive family content and candidate accessory functions

The central result is predictive: protein-family content retains information about *nifHDK* status across held-out genus labels after label-dependent filtering is restricted to training data. Simple genome-size summaries also predict the label, and family-content logistic regression performs better across the tested partitions. The predictive signal therefore depends on which families a genome contains, beyond the broad association between genome size and diazotrophic potential. These results do not isolate causal accessory functions. The two inventories instead provide different starting points for discovery: high carrier purity and positive residual association, with model importance as a separate ordering. Engineering nitrogen fixation still motivates experiments on oxygen management, reductant supply, energy balance, cofactor assembly, transport, stress protection and regulation. Related practical motivations appear in solar-powered oxic N2-fixation and bio-solar nitrogen-economy frameworks [7, 26], while metabolite-based prediction provides a complementary signal ranging from diagnostic to biologically implicated outside genomic and proteomic content [27].

Protein families provide a unit for connecting genome content with protein-function annotation and proteomics. Their utility depends on clustering and mapping quality: a functional name can span multiple sequence clusters, and a cluster-level absence need not imply loss of the named function. The resulting resource is most useful for choosing testable contrasts among candidates and appropriate controls.

### What model comparison and ranking can establish

Using four algorithms shows how prediction depends on the modeling approach, but comparable discrimination does not distinguish if a protein family is additive or necessary for the physiology. Importance can be redistributed among correlated families, and absolute importance does not specify effect direction or causality. Inventory membership is determined by the carrier association, without a consensus-importance threshold. Full-panel and training-fold importance rankings are provided separately. Tier A reflects annotation scores and need not contain the families ranked highest by the classifiers.

Prediction evaluates the modeling procedure, whereas experimental prioritization requires inspection of individual families. Genus-label blocking does not guarantee phylogenetic or sample-identity independence. The assembly-status, grouping and taxonomy sensitivities qualify generalization, while model ranks and study-specific annotations identify questions for follow-up rather than experimentally established requirements.

### Physiological annotations guide experimental selection

The overlays provide different annotations with substantial dependencies. The related atlas compares product labels on largely overlapping genomes. Interpretive labels for oxic nitrogen fixation are distinct from the gene-level predictions in the prior *Anabaena* analysis [20].

Proteomics records study-specific responses of mapped proteins, and the condensate resource supplies predicted sequence-based ranks. The measured dependence of proteomics annotations on detection opportunity, and of the condensate inventory contrast on prevalence, limits what agreement between those layers can establish.

An active-phase-up proteomics annotation identifies a physiological time or cell type in which to test a mapped candidate. Proteins detected in more studies have more opportunities to receive that annotation. The candidate tables therefore retain study-specific timing and the number of mapped studies, so readers can assess detection opportunity when choosing experiments.

The condensate-driver overlay has a similar limitation. The apparent residual-association inventory advantage attenuated after adjustment for carrier-genome prevalence, so the raw contrast does not distinguish accessory-specific protein organization from broader family-distribution effects. Condensate rank can motivate localization and biophysical experiments, while prevalence-matched comparisons and direct measurements remain necessary to test specificity.

### Engineering implications

The two inventories are useful for choosing engineering experiments. The high-purity set supplies high-carrier-purity context, including expected Nif machinery; the residual-association set orders for candidates using model importance. Separating the inventories keeps broadly shared support functions in view as engineering candidates even when they are poor diagnostic markers of diazotrophy. Annotations suggest tests of protein maintenance, nitrogen assimilation, carbon and redox metabolism, electron transfer, and cofactor handling. Carrier representation across genus-based morphotype proxies can help select candidates for portability tests, but neither breadth nor score establishes portability. For established Nif components, failure of a particular cluster to pass a breadth flag warrants a review of sequence membership and representation before that component is excluded from consideration.

Families with limited functional annotation remain candidates for discovery, but Other / unassigned and Novel/uncertain are not reliable synonyms for unknown function. The 74 residual-association families flagged by generic or hypothetical product-name patterns require domain and sequence review before novelty can be assessed. Model rank, carrier distribution and study-specific proteomics can guide that review. Recent nitrogen-responsive extracellular proteomics in *Anabaena* also motivates tests of envelope and extracellular functions, without establishing that they are required for nitrogen fixation [28].

### Protein maintenance and fixed-nitrogen assimilation remain testable hypotheses

The inventory suggests experiments on protein and RNA maintenance during nitrogen-fixing states. The active-phase criterion retains the biological timing of observed abundance changes, but detection opportunity still influences which families receive repeated support. Protein-maintenance hypotheses should therefore be evaluated against candidates matched for detectability and family prevalence, followed by perturbations that distinguish a general growth defect from a nitrogen-fixation-specific effect. One test is to compare candidate perturbations during diazotrophic growth and growth supplied with fixed nitrogen, measuring nitrogenase activity separately from overall growth.

Protein-quality control, RNA regulation, carbon/nitrogen homeostasis and the supply of ATP, reductant and carbon skeletons provide established physiological contexts for the annotated candidates [21, 22, 29–34]. Clp, chaperone, peptide-release, NYN-domain and tRNA-modification families motivate tests of maintenance demands during diazotrophy; amino-acid-biosynthesis and NtcA/PII-associated families motivate tests of nitrogen-assimilation demand. Their representation among high-scoring families does not demonstrate increased demand or a nitrogen-fixation-specific requirement. Those hypotheses require measurements and perturbations that separate general growth effects from effects on nitrogen fixation.

Candidates with high condensate-driver ranks offer a complementary experimental question: whether protein localization or organization changes during the relevant nitrogen-fixing state. Testing this question requires direct localization or biophysical measurements and appropriate prevalence- and detectability-matched comparisons. The current overlay cannot establish selective condensate association of the residual-association inventory.

For families with limited functional annotation, domain architecture, intrinsic disorder, low-complexity sequence, interaction partners, and localization can help choose experiments. These features should be inspected directly rather than inferred from membership in an unassigned keyword bin or from a high composite tier.

### Relationship to microbial association testing

This workflow complements microbial pan-GWAS methods that test family- or variant-level associations and address population structure through different approaches [13–15]. Here, genus-blocked evaluation measures predictive discrimination, while the full-panel importance ranks and annotations guide experimental selection. These results are not calibrated family-level association significance tests.

## Limitations

First, the diazotrophy label is based on genomic *nifHDK* detection and therefore measures diazotrophic potential without verifying nitrogen-fixation activity under a specific condition. Some genomes may carry inactive, cryptic, or conditionally expressed systems, and a Mo-nitrogenase *nifHDK* label leaves alternative nitrogenases unresolved.

Second, stored genus labels do not ensure phylogenetic or sample-identity independence. The cohort and grouping sensitivities retain the original clustering and prevalence-defined vocabulary rather than rebuilding an atlas from reviewed assemblies and raw proteins. Deeper phylogenetic structure, unequal genus representation, unstratified folds and genome-size associations remain. The full-data family ranking is a discovery resource; procedure-level prediction does not independently validate each family.

Third, product annotations and name-based groupings can be incomplete or misleading. Standardized functional re-annotation is needed before comparing category enrichment or inferring biological novelty. The annotation procedures and their distinct roles are specified in Methods and the repository annotation note.

Fourth, morphotype flags use genus-based proxies and contribute substantially to the heuristic score. The related 449-genome morphology table was also generated by genus-name lookup. It covered 416 primary genomes, assigning morphology to 379 and marking 37 unknown; 10 primary genomes were absent. Among the 96 shared, assigned diazotroph genomes, its unicellular/filamentous labels agreed exactly with the primary broad proxy. This is annotation concordance, not independent morphological validation or evidence of portability.

Fifth, the current presence/absence matrix ignores copy number, regulatory context, promoter architecture, operon boundaries, protein abundance, and environmental condition. These layers are likely important for engineering. At 80% clustering identity, proteins with the same functional name can occur in separate families; absence of one cluster does not establish absence of the function. Cross-atlas comparison is additionally limited by incompatible family identifiers and the missing related-atlas membership map. The prioritization score omits transferred atlas terms, while its remaining weights are heuristic rather than empirically calibrated.

Sixth, the proteomics and condensate annotations depend on mapping and measurement choices. Only 2,125 of 31,021 condensate-ranked proteins mapped to 1,311 families, including 804 mappings available only through a cluster-accession fallback. With those fallback mappings omitted and the original 1,311-family rank denominator held fixed, one Tier B family fell below its threshold: acetolactate synthase small subunit GF_02157. The shortlist decreased from 167 to 166 and active-up annotations from 65 to 64; all 22 Tier A families and their best condensate annotations were unchanged. Five ranked rows used WP accessions assigned to more than one family elsewhere in the mapping table; removing those rows did not change shortlist membership. These sensitivities do not establish whether a particular mapping is biologically correct. Condensate rank does not establish phase separation or a mechanistic requirement.

Proteomics sources differ in contrast, normalization and platform selection, including cysteine-targeted heterocyst analysis; their active-up labels are not equivalent statistical effects. The confound models use mapped-study count as a detectability proxy, product-name patterns for functional groups and some small prevalence strata. Matched quantitative measurements, sequence checks and perturbations would permit stronger tests; attenuation of aggregate contrasts does not rule out roles for individual candidates.

## Conclusions

Protein-family content predicts *nifHDK*-defined diazotrophic potential in this cyanobacterial panel. Training-fold filtering, size baselines and dated assembly and taxonomy checks define the scope of that result. The candidate resource separates 981 high-purity families from 473 positive residual associations, with three ubiquitous controls retained for comparison. Model importance and study-specific annotations provide different ways to inspect these families. The heuristic score prioritizes 167 candidates for genetic and physiological tests. Detection opportunity, prevalence and mapping uncertainty limit what annotation agreement can establish about their functions.

## Methods

### Genome selection and diazotrophy labeling

The original pipeline collected cyanobacterial assemblies using NCBI Datasets [35, 36]. Of 489 records, the archived RefSeq-accession (GCF) rule excluded 63; the requirement for complete assemblies excluded none further, yielding the fixed 426-assembly panel. Exact versioned accessions, the matrix and metadata are preserved at recorded repository revision ac2e43a6 (13 January 2026 UTC). This date records archive provenance; the original retrieval timestamp was not captured. Diazotrophy labels were assigned by detecting NifH, NifD and NifK proteins using profile HMM searches with HMMER [37]. The classifier target was the binary diazotrophy label, requiring a best independent domain E-value <1e-50 for each of NifH, NifD and NifK. This rule yields 112 positives; a 1e-30 cutoff gives the same labels in the retained panel. Looser scan-presence flags retained in the metadata were not used as model targets. The supplied profiles were traced to NFixDB [38], with exact profile-content matches after line-ending normalization and matching seed-alignment checksums. Each alignment contains 29 rows; the NifH alignment includes one repeated identical reference, leaving 28 distinct headers. The archived profiles were used without deduplication. Public source files and accession headers are provided in the repository. Original predicted proteomes and domain-hit tables were unavailable for a complete upstream replay.

### Protein retrieval and family construction

The archived workflow retrieves predicted proteins using an NCBI download helper, with Datasets preferred and assembly-summary retrieval as a fallback, and prefixes FASTA headers with assembly accessions. The driver invokes MMseqs2 [17] with minimum identity 0.8, coverage 0.8, greedy set-cover clustering (cluster-mode 0) and bidirectional coverage (cov-mode 0). Cluster assignments are converted into protein-to-family maps. The prevalence filter retains families with at least 40 carrier genomes, producing the fixed 426 × 2,286 binary matrix. The historical MMseqs2 executable version and automatically chosen sensitivity were not recorded in the archived workflow. Prediction analysis starts from the saved family matrix; raw cluster input and execution were not replayed.

### Supervised modeling

The classifiers were regularized logistic regression, random forest [18], gradient boosting, and XGBoost [19], implemented with scikit-learn [39] and xgboost. Training used fixed random seeds and custom genus-level cross-validation. In the primary analysis, the 75 stored genus-name tokens were shuffled with seed 42 and assigned round-robin to folds, keeping each token within a single fold. Folds are unstratified by diazotrophy label. Random forest used 100 trees, max depth 10, and minimum leaf size 5. Gradient boosting used 100 estimators, max depth 5, and learning rate 0.1. Logistic regression used a maximum of 1,000 iterations. XGBoost used 200 estimators, max depth 5, learning rate 0.1, subsample 0.8, colsample_bytree 0.8, and log-loss evaluation. All hyperparameters were held fixed for cross-validation, using scikit-learn 1.6.1 and XGBoost 3.1.1. Metrics were summarized across the same five genus-blocked folds.

Within each training fold, we excluded families whose diazotroph fraction among training carriers was ≤0.10 or ≥0.90 and families represented in fewer than three training genera. Exact integer-count comparisons enforce the boundaries. Neither test labels nor test-genus occurrences enter these filters. Logistic-regression scaling is fitted on training data only. The family vocabulary and ≥40-genome prevalence filter were fixed before evaluation; protein clustering was not repeated within folds. Fold assignments, selected features, probabilities, runtime versions and input hashes are provided in the repository. Importance rankings from training-fold models are reported separately from the full-panel inventory ranking.

We report unweighted means and sample standard deviations of five fold metrics, plus pooled out-of-fold summaries. Average precision uses the stepwise precision-recall definition, not trapezoidal PR area. For genus-weighted summaries, each genome receives weight 1/n_g, where n_g is the number of genomes from its genus, giving every genus equal total weight. Genus-weighted pooled AUC is not a mean of within-genus AUCs. Two single-variable logistic baselines use assembly total ungapped length or the number of present families in the fixed matrix, with training-only scaling and the same folds. These baselines assess broad predictive structure and do not estimate causal effects of genome size.

Partition sensitivity used seeds 42–51 without tuning: the same genus splitter, exact training-fold filters and logistic-regression settings were applied to family content, assembly length, retained-family count, and the separately specified family-plus-length model. We summarize paired differences and ranges without a significance test across overlapping partitions. The seed-42 four-model evaluation remains the primary performance summary.

We retrieved NCBI reports for the exact assembly versions on 30 September 2026 and repeated model fitting for current records, preserving surviving genomes’ original folds. Statuses were not backdated to original collection. Additional status, sample-identity, genus-rank and training-prevalence sensitivities used the fixed family vocabulary and training-only filters without retuning. Their full designs, dated inputs and model-specific results are documented in the accompanying Cohort and taxonomy sensitivity analysis note.

### Candidate inventory construction

Carrier purity is the fraction of a family’s carrier genomes labeled *nifHDK*-positive. The high-purity inventory contains 981 families with carrier purity ≥0.90. After excluding high-purity and comparison-pure families, the residual-association inventory contains 473 families with a positive difference between P(*nifHDK*-positive | family present) and P(*nifHDK*-positive | family absent), computed across the full panel. Three ubiquitous controls remain in the 476-entry inventory: cytochrome b6 GF_01297, ribosomal S10 GF_01899 and RpaB GF_01945. Their contrasts cannot be estimated because every genome is a carrier. The source implementation included them by substituting zero for the empty non-carrier group; they are excluded from shortlist eligibility. All 476 entries appear in the four model-importance tables. Mean within-model percentiles of absolute importance determine their order, without a significance, stability or nonzero-importance threshold. The source data labels Model-Supported and Highly Pure correspond to the residual-association and high-purity inventories, respectively. These identifiers remain in machine-readable files and the workbook; the repository terminology guide maps them to manuscript terms. Prediction evaluation uses separate training-fold filters and does not redefine the full-panel inventories.

### Genus-based morphotype representation

Candidate families were further annotated with morphotype-breadth flags using genus-based proxies because the current metadata lacked a curated morphology column. The strict unicellular proxy included diazotroph-labeled genomes with the stored genus-label tokens “Candidatus”, *Crocosphaera*, *Euhalothece*, *Gloeothece*, *Halothece*, *Rippkaea*, *Synechococcus*, *Synechocystis*, and “cyanobacterium”. The broad unicellular proxy added *Chroococcidiopsis*, *Cyanobacterium*, and *Pleurocapsa*. For each proxy, all remaining diazotroph-labeled genomes were treated as a filamentous or non-unicellular comparison proxy.

The primary morphotype bridge requires at least one broad-proxy unicellular diazotroph carrier and at least one filamentous-proxy diazotroph carrier. Additional criteria require at least three broad- or strict-proxy unicellular carriers, with at least one or at least three filamentous-proxy carriers. These criteria describe carrier representation under the genus-based assignments. A separate 12-family set, defined by *nif*/nitrogenase keywords, supplies reference annotations and scoring exclusions. We retain its membership for reproducibility while distinguishing established structural Nif components from other name-based entries.

### Functional annotation schemes

Family product annotations were derived from representative proteins and modal product strings where available. Product-name keywords define coarse functional categories; a separate workbook grouping records broader functional context. Interpretive labels distinguish plausible support for oxic nitrogen fixation, housekeeping or lineage context, and uncertain function.

These labels are distinct from the prior gene-level prediction model and contribute no composite-score term. Regular expressions identify generic or hypothetical product names and matches to a broad proteostasis, translation and amino-acid-metabolism pattern. Such matches do not establish curated functional classes or unknown function. The housekeeping/lineage indicator used for score penalties and tier exclusions is computed from product strings and the keyword categories. The repository data dictionary specifies the column names, rules and diagnostic outputs. Figure 3 uses separate product-annotation groups for display; those groups do not change categories used in scoring, housekeeping indicators or composite scores.

### Protein-family versus gene-level comparison

A separate gene-level top-100 list from genus-blocked random forests on gene-level features was mapped into the protein-family space. The main comparison reported here used all genes available for each gene-driven feature family and recorded whether each feature mapped to the residual-association or high-purity candidate inventories. Functional composition of the gene-level top-100 list was compared with the protein-driven top-100 list using conservative keyword bins. The 200-row category table and plotting inputs reproduce the reported category totals. The eight-bin diazotrophy-physiology aggregate follows the archived README; category assignments remain keyword-based.

### Related-atlas concordance and proteomics prioritization

The related-atlas comparison uses the 449-genome analysis in the ERISE-BNERC repository identified in Results. We compared normalized product/function labels because family identifiers differ between clustering runs. Generic labels such as hypothetical, uncharacterized, DUF-containing, domain-containing-protein and broad family-protein names were tracked separately from informative product matches. Matches were then intersected with the 1,457-family candidate inventory, its residual-association and high-purity subsets, and the active-phase-up proteomics annotations. Because the two atlases share 416 genomes, this is a consistency check on largely overlapping data. Neither raw-identifier overlap nor product-label agreement contributes to the prioritization score. Family-level transfer requires a protein-accession or sequence-based crosswalk. The Related-atlas comparison note documents source versions, reported model metrics and scoring provenance.

External proteomics was used as a prioritization layer. Proteins from the source studies were mapped to the current protein-family namespace through protein-accession and locus-tag crosswalks, including legacy identifiers and current WP accessions, without product-name matching. Positive proteomics score required increased abundance in the organism-appropriate nitrogen-fixation-active state or cell type: dark nitrate-free *Crocosphaera*, daytime *Trichodesmium*, late-light/early-dark PCC 7822, or heterocyst-enriched *Nostoc* fractions.

Active-phase-down proteins, transition-window dynamics, broad nitrogen responses, and non-target-phase responses were retained as annotations but did not add score. The resulting priority table preserved each evidence type as a separate column: current family ID, current model status, active-phase-up proteomics support, condensate-driver rank, morphotype-breadth fields, and heuristic functional interpretation.

The condensate-driver resource supplied an additional annotation layer. Its 31,021 rows mapped to 1,311 current families through 1,321 direct protein-accession matches and 804 cluster-accession fallback matches. Direct accession matches took precedence, including 22 rows for which the two routes disagreed. No mapped row was labeled a training positive in the supplied resource; this check does not resolve all possible dependence on the ranking model. The best mapped score and rank summarized each family. Unadjusted enrichment summaries used one-sided Mann-Whitney tests for scores and Fisher exact tests for upper rank fractions; prevalence-adjusted diagnostics are described below. For fallback-mapping sensitivity, affected families were placed using their best retained direct score and rank against the original family reference, excluding their own original entry and retaining the denominator of 1,311; unaffected percentiles stayed fixed. A second conservative analysis set a family’s condensate component to zero when its original best annotation was lost. Both rules gave the same sole tier migration. We also removed rows using WP identifiers assigned to multiple families. Assembly-plus-WP keys were unambiguous in the retained map, and independent reconstruction of the proteomics assembly-specific joins reproduced all 6,426 saved mapped-family sets for source protein/study records. This check validates those joins against the supplied map, not the biological identity of every historical protein crosswalk.

### Evidence-layer mapping and sensitivity summaries

The four literature sources use study-specific response rules. For Panda, active-up requires a nitrate main-effect or interaction ANOVA p value ≤0.05 and dark nitrate-free/nitrate-replete logFC ≥0.25; a dark-specific pairwise significance test is not required. For Sandh, active-up requires a heterocyst/filament log2 ratio ≥1 in the current-study comparison. For Welkie, the range across four supplied CT-normalized values must be ≥1 in those reported units, with the maximum at D0 or D3. For Held, (maximum − minimum)/mean abundance must be ≥0.5, with the maximum at 6–15 hours after dawn. The latter two are temporal proxies without a per-protein significance requirement. Repeated records count at most once per study and family.

Active-down, transition-window and other responses remain non-scoring annotations. Source-table checks verified that rounded Held metrics and peak ties do not alter the retained classifications.

### Composite prioritization score and tier definitions

The composite score is a heuristic for choosing experimental candidates. Inventory membership contributes M = 2.0 for residual-association families and 1.0 for high-purity families. Classifier rank does not enter this component. Proteomics, condensate rank and morphotype representation supply the remaining positive components.

Proteomics contributes P = min(u, 3) + b, where u counts studies with an active-up annotation. The context bonus b is 0 for zero or one context, 0.5 for two, and 1.0 for three. Context groups are unicellular (*Crocosphaera* and PCC 7822), heterocyst (*Nostoc*), and nonheterocystous filament (*Trichodesmium*). Condensate score C is 0.2, 0.3, 0.4 or 0.5 for the highest applicable top 20%, 10%, 5% or 1% band, respectively, and zero otherwise.

Morphotype score B is 1.25 for the primary bridge plus 1.5 for strict unicellular breadth. Penalties H sum 1.0 for the housekeeping/lineage indicator and 1.0 for membership in the *nif*/nitrogenase keyword set. The adjusted score is M + P + C + B − H + I_HP, where I_HP = 1 for high-purity families and 0 otherwise. This inventory offset is part of the baseline weighting, not additional external evidence. Scores are rounded to three decimal places; missing scored annotations contribute zero. The weights and thresholds were fixed before the 167-family shortlist was calculated.

Ubiquitous controls, members of the *nif*/nitrogenase keyword set and families with the housekeeping/lineage indicator are excluded before tier assignment. The controls retain their numeric annotations and scores in the complete table. Among eligible families, Tier A requires residual-association membership and an adjusted score ≥6.0; Tier B comprises the remaining residual-association families with scores ≥4.5. The high-purity tier requires high-purity membership and a score ≥4.5. These bands contain 22, 140 and 5 families, respectively.

Additional rules allow Tier C for other families scoring ≥4.5, Tier D for remaining families with at least two active-up studies, and Tier E otherwise; no C or D families occur here. Figure 3 groups Tier A families by product annotation and orders them within groups by decreasing carrier purity, then family ID. Display order does not affect selection. Complete selection fields accompany the shortlist. Machine-readable tier identifiers, including Tier HP for the high-purity tier, are defined in the terminology guide.

### Diagnostics of annotation dependence

We examined annotation dependence across the 1,457-family inventory. These diagnostic populations retain the three ubiquitous controls, whereas shortlist eligibility excludes them; each summary states its population. Among residual-association families mapped in at least one proteomics study, logistic regression modeled any active-up annotation against mapped-study count, strict unicellular breadth and full-panel model-importance percentile. Confidence intervals are model-based Wald intervals and do not adjust for dependence among families. We compared mean per-family active-up/mapped-study ratios by breadth indicator. For condensate-mapped candidates, logistic regression modeled top-10% rank membership against inventory and log(mapped-protein count), then added log(carrier-genome count). One-sided Fisher tests within carrier strata were exploratory and uncorrected for multiple testing. The repository provides the diagnostic and regression scripts.

We calculated score correlations and product-name-pattern summaries for the stated tiers. The repository annotation note provides the category co-occurrence methods and full results. These diagnostics describe dependence among annotations; they do not test individual family functions.

### Use of large language models

AI-assisted tools supported manuscript drafting and editing and the preparation and checking of analysis code, diagnostic summaries and figures. Authors have reviewed and approved of these materials, and take responsibility for the study design, analyses, results, interpretations and conclusions.

## Declarations

### Ethics approval and consent to participate

Not applicable.

### Consent for publication

Not applicable.

### Availability of data and materials

The analysis files are available on github (https://github.com/jamesyoung93/Diazotrophic_Cyanobacteria_Pangenome/tree/main/manuscri pt_release_2026_10). The release contains exact assembly versions and dated NCBI status and taxonomy snapshots, the fixed protein-family matrix, candidate inventories, scores, saved predictions, analysis scripts, runtime manifests, source-study checks and figure inputs.

Repository methods notes document cohort sensitivity, annotation rules, the related-atlas comparison and terminology. The candidate workbook is annotations/results/current167/candidate_families.xlsx within the release.

### Competing interests

The authors declare that they have no competing interests.

### Funding

This material is based upon work supported by the U.S. National Science Foundation under NSF EPSCoR Collaborative Research-E-RISE RII: BioNitrogen Economy Research Center Award No. <u>2416911</u> and partially supported by the South Dakota Agricultural Experiment Station– USDA Hatch projects and NIFA Award No. <u>2023-67022-39594</u>.

### Authors’ contributions

James Young, Ruanbao Zhou, Etienne Z. Gnimpieba, and Shiva Aryal conceived and coordinated the study. James Young led the project including bioinformatic and machine learning methodology, integrated the analytical results, incorporated proteomic validation datasets, performed biological interpretation, prepared figures, and wrote the original manuscript draft. Shiva Aryal developed and performed complementary machine-learning and comparative-genomics analyses, generated supporting scripts and validation analyses, curated analytical datasets, contributed visualizations, and reviewed and edited the manuscript. Dillon Nelson contributed to study development, data interpretation, and manuscript review. Dheeraj Raya contributed to manuscript development, scientific discussion, writing, review, and revision of the manuscript. Etienne Z. Gnimpieba provided scientific leadership in workflow development, machine-learning design, study interpretation, manuscript revision, and supervision. Ruanbao Zhou contributed to study design, manuscript revision, supervision, and project oversight. All authors reviewed and approved the final manuscript prior to submission.

## Acknowledgements

This research was supported in part by South Dakota State University’s High Performance Computing resources.

## Notes

https://github.com/jamesyoung93/Diazotrophic_Cyanobacteria_Pangenome

### Competing Interest Statement

The authors have declared no competing interest.

